# Potentialities of biotechnological recovery of hydrogen and short- and medium-chain organic acids from the co-fermentation of cheese whey and Yerba Mate (*Ilex paraguariensis*) waste

**DOI:** 10.1101/2021.07.16.452613

**Authors:** Antônio Djalma Nunes Ferraz Júnior, Laura Fuentes, Victoria de la Sovera, Patricia Bovio-Winkler, Felipe Eng, Mariángeles Garcia, Claudia Etchebehere

**Affiliations:** Laboratorio de Ecología Microbiana, Bioquímica y Genómica Microbiana, Instituto de Investigaciones Biológicas “Clemente Estable”. Av. Italia 3318, Montevideo, Uruguay; Laboratório de Processos Biológicos, Escola de Engenharia de São Carlos, Universidade de São Paulo (LPB/EESC/USP). Av. João Dagnone 1100 - Santa Angelina, 13563-120, São Carlos, SP, Brasil

**Keywords:** Central Composite Design, Response Surface Methodology, Butyrate-type fermentation, Caproate-type fermentation, Lactate-type fermentation

## Abstract

Co-fermentation of cheese whey (CW) and thermal-alkaline pre-treated Yerba Mate (*Ilex paraguariensis*) waste (YMW) was performed aiming to produce biohydrogen and/or short- and medium-chain organic acids. Central Composite Designs (CCD) was chosen as the experimental design for evaluating the combinations of three independent variables namely YMW concentration, pH and inoculum concentration in hydrogen yield (H_2_Y; response variable). The increase of inoculum and YMW concentrations had positive effect in biohydrogen production and yield (H_2_Y_max_ of 1.35 mMH_2_.g^-1^ VS _added_) whereas the initial pH had no significant effect on it. Hydrogen was produced as a coproduct to butyrate mainly. Acetate from homoacetogenesis was accounted in all conditions evaluated. The CCD also indicated operating conditions to produce moderate-to-high concentrations of short and medium-chain organic acids such as butyrate (~135 mM), caproate (~45 mM) and lactate (~140 mM). 16S rRNA gene sequences analysis revealed five groups of microorganisms related to hydrogen, lactate and caproate production, ethanol-hydrogen co-production and hydrogen consumption.

**Highlights:**

- Co-fermentation improved hydrogen production in up 7.5-folds compared to the sole CW-fed system.
- The initial pH had no effect on hydrogen-producing batch reactors.
- Hydrogen was produced as a coproduct to butyrate.
- Design of experiment indicated operating conditions to the production of lactate and caproate.

## 1. Introduction

Fermentation is one the first steps of residues decomposition by microorganisms. In this step, hydrogen is the most desired coproduct for being a versatile energy carrier used for fossil fuel refining and production of chemicals, including biofuels (IEA, 2019). Other coproducts of industrial interests such as short- and medium-chain organic acids, alcohols and solvents can also be obtained through fermentation process which makes it an ideal multipurpose technology (Borin et al., 2019; Luongo et al., 2019; Mota et al., 2018).

In general terms, fermentation technology comprises a cascade of reactions which are primarily related to the production and consumption of hydrogen, considering a fermentative system using non-sterile mixed cultures (Levin et al., 2004). In that case, high production of hydrogen is associated to the mixture of acetate and butyrate fermentation route end-products whilst its low production is associated to other reduced end-products such as acetone, butanol, ethanol and lactate (Ferraz Júnior, 2013). Finally, hydrogen consumption is reported in methanogenesis and homoacetogenesis routes. These reactions will be reported in depth in *section 3.6*.

The success of fermentative systems is allied to multiple factors (substrate-type, temperature, pH, inoculum, regime operation) which are interrelated (Akhlaghi et al., 2017; Koyama et al., 2016). The operating pH (controlled along the process) is an important individual-factor in fermentative systems. It indicates the hydrolysis and fermentation degree; determines the activity of hydrogenase and the metabolic routes (Kim et al., 2011). Extreme high pH values can negatively affect the activity of hydrogen-producing microorganisms as well as extreme low pH values can result in inhibition of the hydrogenase activity (Mohd Yasin et al., 2011) diverting the corresponded pathways to the production of solvents (i.e., alcohols) (Fuess et al., 2018). Similarly, the relative concentration of active biomass (inoculum) in the system and the substrate available to be consumed is expressed as food-to-microorganisms (F/M) ratio. This ratio can shift from substrate-limited to substrate-sufficient growth but also to substrate-excess unbalancing the anabolic and catabolic reactions and, thus, affecting the yield of substrate conversion into by-products (Akhlaghi et al., 2017; Liu, 1996).

Different wastes and wastewaters have been used as feedstock in fermentative systems including agricultural and food industry wastewater, lignocellulosic biomass and organic fraction of municipal solid waste (Castelló et al., 2020). However, some feedstock can present undesirable features or even a nutritional deficit which may affect the process to proceed properly. For instance, algal biomass has been adopted as the main carbon source for sewage sludge fermentation in order to dilute the inherent inhibitors to the latter residue (Yin et al., 2021). Similarly, rich-protein substrate (microalgae) and rich-carbohydrate substrate (macroalgae and rice residues) have been used as mixed substrate to achieve better ratios of carbon-nitrogen and improve the performance of fermentative organic acids and hydrogen production (Sun et al., 2018; Xia et al., 2016). The simultaneous fermentation of two or more residues, also known as co-fermentation, might represent an alternative to mitigate the aforementioned drawbacks (Grosser and Neczaj, 2018; Yang et al., 2019) and increase the production of target products.

Cheese whey (CW) is a residual nutrient-rich liquid stream from dairies industries that has been extensively studied to fermentative purposes (Basak et al., 2018; Lovato et al., 2018; Rao and Basak, 2021). However, hydrogen production instability and process inhibition by accumulation of organic acids have been reported and attributed to its lack of alkalinity (low pH) and high organic matter concentration, respectively (Fernández et al., 2014; Lovato et al., 2018, Lovato et al., 2021).

Yerba Mate (*Ilex paraguariensis*) waste (YMW) is one the most important lignocellulosic residue in Southern Cone of Latin America (Argentina, Brazil, Chile, Paraguay, and Uruguay) and after its thermal-alkaline pretreatment might be a co-substrate for CW fermentation able to increase bioproducts production. The thermal-alkaline pretreated YMW presents a high pH (13.1 ± 0.6) (Ferraz Júnior et al., 2020) and might be able to “buffer” the system by increasing the pH to suitable values of fermentative process without additional costs with alkalis.

Co-fermentation of CW and YMW for biohydrogen and/or short- and medium-chain organic acids production has not been described in any literature before, therefore, it represents a novelty and the aim of this study. The design of experiments (DoE) was used as a systematic method to investigate fundamentals factors of the process (initial pH, concentration of inoculum and YMW) in batch-mode to attain the production of hydrogen and/or short- and medium-chain organic acids. High-throughput sequencing (HTS) technology was also performed to assess the microbial community dominant in the co-fermenting system.

## 2. Material and methods

### 2.1. Substrates: Cheese Whey (CW - substrate) and Yerba Mate Waste (YMW – co-substrate)

Cheese Whey (CW) was collected from an artisanal cheese producer (daily milk production of 6000 – 9000 L.d^-1^) in Uruguay. Yerba Mate Waste (YMW) was used as a co-substrate. Briefly, YMW was thermal-alkaline pretreated to unlock the carbohydrates/sugar prior to co-fermentation. The composition of substrate and co-substrate are depicted in Table 1. Details about YMW generation and the pretreatment conditions are presented in Ferraz Júnior et al. (2020).

**Table 1.** Characterization of cheese whey (main substrate) and thermal-alkaline pretreated Yerba Mate waste (co-substrate).

| Variable | Unit | CW | Thermal-alkaline pretreated YMW |
| --- | --- | --- | --- |
| pH | - | 4.1±0.8 | 13.1±0.6 |
| Total reducing sugars (TRS) | g.L<sup>-1</sup> | 21.5 | 42.7 |
| Total chemical oxygen demand (TCOD) | g-O<sub>2</sub>.L<sup>-1</sup> | 50±2.5 | 135.0±6.9 |
| Soluble chemical oxygen demand (SCOD) | g-O<sub>2</sub>.L<sup>-1</sup> | 31.5±0.4 | 98.0±4.6 |
| Total solids (TS) | gTS.L<sup>-1</sup> | 33.2±0.6 | 101.7±2.8 |
| Total volatile solids (TVS) | gTVS.L<sup>-1</sup> | 31.0±0.8 | 50.2±3.3 |

### 2.2. Inoculum

Organic compost was used as inoculum (T.Res.Or, Montevideo, Uruguay). According to the manufacturer’s information the compost presents a pH of 7.2, the moisturize content of 25.1%, and TVS of 35%.

### 2.3. Box-Wilson Central Composite Design (CCD – 2^k^) with center point repetition

Central Composite Designs (CCD) was chosen as the experimental design for evaluating all combinations of all factor-levels of each factor (Box G.E, Wilson, 1951) thus, enabling the estimation of all factors and their interactions in the process of biohydrogen production from co-fermentation of CW and YMW. Design levels were determined. Three independent variables, namely YMW concentration (X_1_), pH (X_2_) and inoculum concentration (X_3_) were studied resulting in five levels: CCD (±1), center point (0) and axial points (±α) with 3 repetitions at the center point. The values were assumed based on practical values of solid waste management, reports of biohydrogen production with and without pH control (Bina et al., 2019; Koyama et al., 2016; Mota et al., 2018) and practical values of reactors inoculation in relation to its working volume (Table 2). The number of experiments performed was given by Equation 1. Axial points were given by Equation 2.

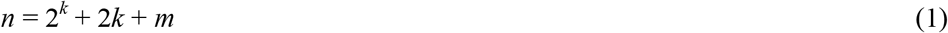

**Table 2.** Three-variables design matrix for biohydrogen production from CW and YMW. Replications of center point or control are in bold.

| S. no. <sup>a</sup> | Coded level |  |  | Uncoded level |  |  |
| --- | --- | --- | --- | --- | --- | --- |
|  | X<sub>1</sub> | X<sub>2</sub> | X<sub>3</sub> | YMW <sup>b</sup> | pH | Inoculum <sup>c</sup> |
| 1 | -1 | -1 | -1 | 5.1 | 5.9 | 10.0 |
| 2 | 1 | -1 | -1 | 20.0 | 5.9 | 10.0 |
| 3 | -1 | 1 | -1 | 5.1 | 11.0 | 10.0 |
| 4 | 1 | 1 | -1 | 20.0 | 11.0 | 10.0 |
| 5 | -1 | -1 | 1 | 5.1 | 5.9 | 25.0 |
| 6 | 1 | -1 | 1 | 20.0 | 5.9 | 25.0 |
| 7 | -1 | 1 | 1 | 5.1 | 11.0 | 25.0 |
| 8 | 1 | 1 | 1 | 20.0 | 11.0 | 25.0 |
| **9** | 0 | 0 | 0 | 12.5 | 8.5 | 17.5 |
| **10** | 0 | 0 | 0 | 12.5 | 8.5 | 17.5 |
| **11** | 0 | 0 | 0 | 12.5 | 8.5 | 17.5 |
| 12 | -α | 0 | 0 | 0.0 | 8.5 | 17.5 |
| 13 | $\alpha$ | 0 | 0 | 25.1 | 8.5 | 17.5 |
| 14 | 0 | -α | 0 | 12.5 | 4.2 | 17.5 |
| 15 | 0 | $\alpha$ | 0 | 12.5 | 12.7 | 17.5 |
| 16 | 0 | 0 | -α | 12.5 | 8.5 | 4.9 |
| 17 | 0 | 0 | $\alpha$ | 12.5 | 8.5 | 30.1 |

a. The experiments were performed in a random order.
b. Alkaline pretreated Yerba Mate Waste (YMW) in % w/w, as co-substrate.
c. Inoculum, in % w/v (working volume of the reactor).

Where *n* is the number of experiments, *k* is the number of variables and *m* is the number of replicates of the center point by “genuine repetition”.

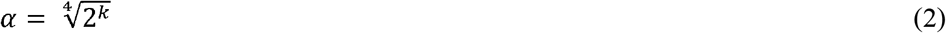

Equation 3 was used to decode α value to access the experimental values of the variables to be studied.

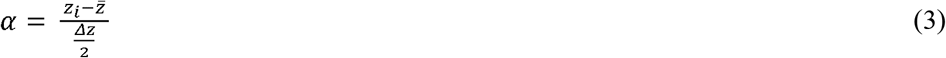

Where α is the coded value of axial point, *z_i_* is the experimental value of the level, 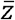 is the average between the lower (-) and higher (+) value of the level which is exactly the value of level zero (0) and Δ*z* is the difference between the lower (-) and higher (+).

The coefficients were obtained using the method of least squares. Linear models were used to evaluate the influence of all the experimental variables of interest and the interaction effects on the response, according to Equation 4.

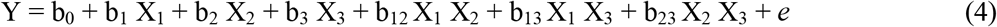

Where Y is the predicted response (hydrogen yield – H_2_Y, in MmH_2_.g^-1^ VS _added_); b_0_ is a constant (average value average of all observations); b_1_, b_2_ and b_3_ are the linear coefficients for the dependent variables (X_1_, X_2_ and X_3_, respectively); X_1_, X_2_ and X_3_ are the variables (YMW concentration, pH and inoculum concentration, respectively); b_12_, b_13_ and b_23_, are the coefficients for the interactions X_12_, X_13_ and X_23_, respectively; and *e* is the random error associated with the model.

The evaluation of significant effects and coefficients was based on statistical decision using analysis of variance (ANOVA) and the Student’s distribution with p-value of 0.05. The significant factors were selected and a response surface methodology (RSM) was used to predict the optimal region on the surface defined by the factors. In this case, the best values of the variables able to produce the higher amount of biohydrogen.

### 2.4. Co-fermentation process of CW and YMW (batch mode)

Co-fermentation of CW and YMW was performed in batch mode. The experiments were performed in parallel according to Table 2. The mixture of feedstocks was performed at room temperature (20-22°C). Schott bottles (DURAN^®^ containing a total volume of 500 mL) were flushed with nitrogen gas, sealed with butyl rubber stoppers, and incubated at 37 °C until hydrogen production ceased. Continuous stirring was kept at 150 rpm. The reaction volume was 200 mL. The pH of all experiments was not controlled during co-fermentation process. Accumulated hydrogen production was measured with a gas volume meter (AMPTS II from Bioprocess Control) previously washed in NaOH solution (12% w/v). Gas production is expressed at standard temperature (0 °C), pressure (1 atm), and zero water–vapor pressure.

### 2.5. Chemical analysis

The pH value was measured by using a pH meter (OAKTON pH 11 series). Total reducing sugars (TRS) were determined using the 3,5-dinitrosalicylic acid (DNS) method (Miller, 1959). Chemical Demand of Oxygen (COD), total solids (TS) and total volatile solids (TVS) were determined according to APHA (2005). Organic acids (C2-C6) were determined by Gas Chromatography equipped with a Flame Ionization Detector (GC/FID) (Adorno et al., 2014). Lactic acid was determined by spectrometry according to Borshchevskaya et al. (Borshchevskaya et al., 2016). Hydrogen (H_2_), Carbon dioxide (CO_2_) and methane (CH_4_) were measured using a gas chromatograph (GC-2014, Shimadzu), equipped with a thermal conductivity detector. A packed column was used with the following dimensions 2 m × 1 mm × 1/16 inch. Temperatures of the injection port and the detector were 120 °C. The initial temperature of the oven was 30 °C, and the final temperature of the column was 110 °C with a temperature increase of 35 °C/min. Ar was used as a carrier gas with a pressure of 8 bar.

### 2.6. DNA extraction, PCR amplification and High-Throughput Sequencing (HTS) of co-fermentation systems samples

Biomass samples were collected from each batch reactor at the end of its operation. However, a composed sample (1:1:1) was generated for the “S.no.” 9, 10 and 11 (replication). 10 mL of samples were centrifuge to separate the biomass (3,000 rpm, 10 min) and genomic DNA was extracted with the ZR Soil Microbe DNA MiniPrepTM kit (Zymo Research) following the manufacturer’s instructions. DNA encoding the 16S rRNA gene was amplified by PCR with primers for the bacteria domain: 520F (5-AYTGGGYDTAAAGNG-3’) and 802R (TACNNGGGTATCTAATCC) (Claesson et al., 2009). Barcodes (10 bp) were added to the amplified 16S rRNA in order to identify the samples after sequencing. The reaction was performed using 1.5 μl of amplified 16S rRNA, 0.5 μl of primers and 12.5 μl of buffer ranger mix (1.5 mM) for a final reaction volume of 25 μl per sample. The conditions were as follow: initial denaturation (95°C for 5 min), 35 cycles of denaturation (94°C for 30 s), hybridization (55°C for 30 s), extension (72°C for 1 min) and final extension (72°C at 10 min). The tagged amplification was purified using the Zymoclean™ Gel DNA Recovery kit following the manufacturer’s protocol. The purified products (tagged amplicons) were sequenced by Ion Torrent PGM technology at Biological Research Institute “Clemente Estable”, Montevideo, Uruguay. The raw reads generated were processed using QIIME software version 1.9.1 (Caporaso et al., 2012, 2011). Low quality reads (coefficient greater than 25) were filtered, trimmed primers, adapters, and barcodes, and reads less than 200 bases in length were eliminated. Chimeras and noise in the sequencing reads were removed leaving high quality reads for the samples. Sequences were clustered into operational taxonomic units (OTU) using UClust algorithm (Edgar et al., 2011), based on the 97% identity threshold (de novo-based OTU picking strategy). OTUs represented by one sequence (singletons) were removed from the analysis. Silva database (version 132) was used for the taxonomic classification of the readings with a confidence threshold of 80%. The raw data was deposited at National Center for Biotechnology Information (NCBI) under accessing number: <u>PRJNA684595</u>.

### 2.7. Calculations and kinetics analysis

The volume of substrate, co-substrate, inoculum and the food/microorganism ratio (F/M) were calculated based on the following system of equations (6) and (7):

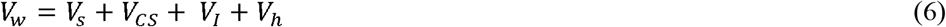

Where, *V_t_* is the working or reactional volume (mL), *V_s_* is the substrate volume (mL), *V_CS_* is the co-substrate volume (mL), *V_I_* is the volume of inoculum (mL) and *V_h_* is the volume of headspace.

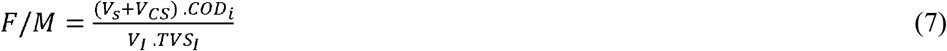

Where, F/M is commonly given in g-COD.g^-1^TVS although is expressed as g-O_2_.L^-1^. *COD_i_* is the initial COD, *TVS_I_* is the total volatile solids of inoculum, in g-TVS.kg^-1^. The dry apparent specific weight (γ_d_) assumed was 600 kg.m^3^.

The experimental data from the optimum condition (Section 2.4) was adjusted to the modified Gompertz equation (GM) using the software package Statistica^®^ 8.0 in order to evaluate the kinetics of the co-fermentation process (Equation 8).

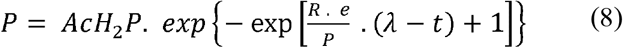

Where, AcH_2_P is cumulative hydrogen production expressed in mM, *λ* is lag-phase time in d, P is hydrogen production potential also in mL, R is the hydrogen production rate in mM.d^-1^ and *e* is exp(l) (i.e., Euler number: 2.71828).

The theoretical expected hydrogen production and the acetate produced from homoacetogenesis were calculated using Equations (9) and (10) as presented in Ferraz Júnior et al., 2020.

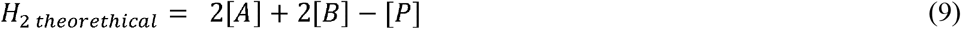

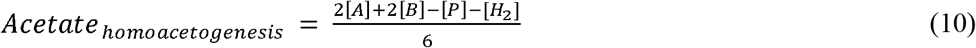

Where, [A], [B], [P] and [H_2_] are the measured acetic, butyric and propionic acids, and the hydrogen concentrations in mM, respectively.

Principal component analysis (PCA) was performed using STATISCA 10 previously described in (Ferraz Júnior et al., 2020).

## 3. Results and discussion

### 3.1. Hydrogen yield: variable response

Full factorial CCD was employed to determine the individual and interactive effects of thermal-alkaline pretreated Yerba Mate (*Ilex paraguariensis*) waste concentration (YMW, % w/w), pH and inoculum concentration (% w/w) on the co-fermentation process which presented cheese whey (CW) as the main substrate. Table 3 presents the experimental (Y) and predicted response (Ŷ) expressed as hydrogen yield (H_2_Y).

**Table 3.** Main results from the CCD experiments. Experimental (Y) and predicted (*Ŷ*) values for the hydrogen yield (H_2_Y, in mM-H_2_.g^-1^ VS _added_) at 5% of significance. Replications of center point or control are in bold.

| S. no. <sup>a</sup> | Uncoded level | | | $H_2Y$ ($mM-H_2 \cdot g^{-1} VS_{added}$) | | Final pH |
| --- | --- | --- | --- | --- | --- | --- |
| | YMW <sup>b</sup> | pH | Inoculum <sup>c</sup> | $\bar{Y}^d$ | $\hat{Y}^e$ | |
| 1 | 5.1 | 5.9 | 10.0 | 0.36 | 0.40 | 3.4 |
| 2 | 20.0 | 5.9 | 10.0 | 0.58 | 0.58 | 3.4 |
| 3 | 5.1 | 11.0 | 10.0 | 0.72 | 0.69 | 4.5 |
| 4 | 20.0 | 11.0 | 10.0 | 0.63 | 0.57 | 3.7 |
| 5 | 5.1 | 5.9 | 25.0 | 0.58 | 0.55 | 4.9 |
| 6 | 20.0 | 5.9 | 25.0 | 1.35 | 1.28 | 4.5 |
| 7 | 5.1 | 11.0 | 25.0 | 0.84 | 0.75 | 4.9 |
| 8 | 20.0 | 11.0 | 25.0 | 1.30 | 1.17 | 5.2 |
| **9** | **12.6** | **8.5** | **17.5** | **0.95** | **0.89** | 4.1 |
| **10** | **12.6** | **8.5** | **17.5** | **0.87** | **0.89** | 4.2 |
| **11** | **12.6** | **8.5** | **17.5** | **0.88** | **0.89** | 4.1 |
| 12 | 0.0 | 8.5 | 17.5 | 0.31 | 0.33 | 3.5 |
| 13 | 25.1 | 8.5 | 17.5 | 0.73 | 0.84 | 4.9 |
| 14 | 12.6 | 4.2 | 17.5 | 0.98 | 0.97 | 3.6 |
| 15 | 12.6 | 12.7 | 17.5 | 0.99 | 1.13 | 5.3 |
| 16 | 12.6 | 8.5 | 4.9 | 0.33 | 0.32 | 3.5 |
| 17 | 12.6 | 8.5 | 30.1 | 0.81 | 0.95 | 4.9 |

a. Experiments were performed randomly. b. Alkaline pretreated Yerba Mate Waste (YMW) in % w/w, as co-substrate. c. Inoculum, in % w/v (working volume of the reactor). d. Experimental value. e. Predicted value.

The different conditions evaluated had a strong influence on hydrogen yield. The highest H_2_Y (1.35 mMH_2_.g^-1^ VS _added_) were obtained when the concentration of YMW and inoculum were at their higher levels and the pH at its lower. In contrast, the lowest corresponding value (H_2_Y; 0.31 mMH_2_.g^-1^ VS _added_) were achieved in absence of YMW and at the lowest concentration of inoculum, indicating that the co-fermentation process was able to increase biohydrogen outputs.

The maximum H_2_Y obtained in the CCD experiments is comparable with data found by other researchers (Table 4). Lee et al., (2008) reported much lower H_2_Y (0.44□mMH_2_.g^-1^ VS _added_) in batch reactors fed with kitchen vegetable wastes. In another study, Dareioti et al., (2014) observed a H_2_Y of 1.06□MmH_2_,g^-1^ VS _added_ from the co-fermentation process of olive mill wastewater, cheese whey and cow manure. In turn, Lucas et al., (2015) evaluated the potential to produce hydrogen from cassava starch, dairy and citrus wastes reaching H_2_Y value of 1.27□MmH_2_.g^-1^ VS _added_ which is similar to the obtained in this study. Exceptionally, Marone et al., (2015) and Basak et al., (2018) reported H_2_Y values four-times higher (4.98-5.69 MmH_2_.g^-1^ VS _added_, respectively) than the maximum value observed in this study. The mentioned authors performed an optimization of substrate composition and kinetics studies for hydrogen production from the co-fermentation of agro-industrial residues with cheese whey as common substrate.

**Table 4.**
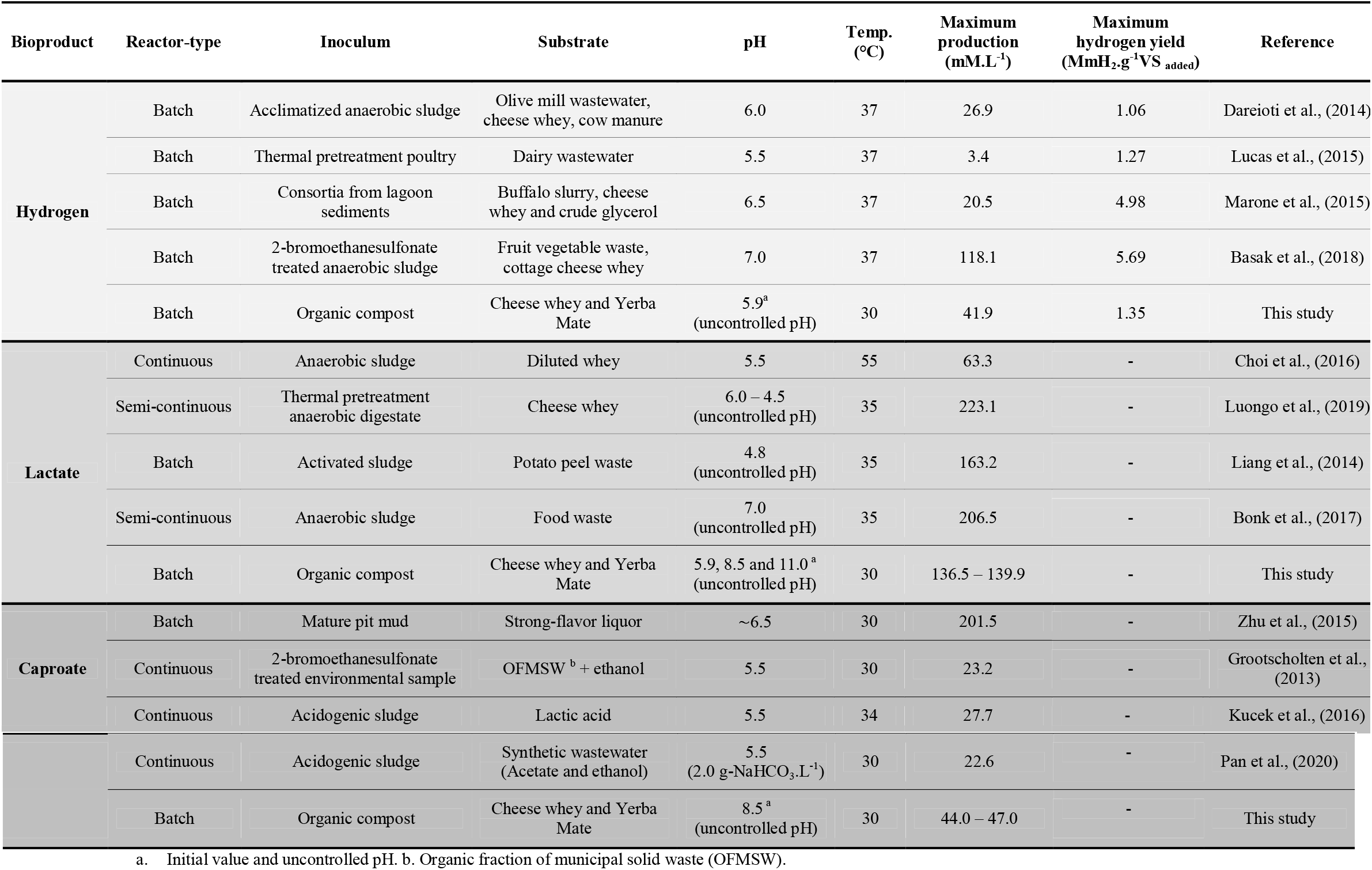
Maximum production and other parameters of BioH_2_, lactate and caproate production by mixed microbial cultures grown on different organic wastes.

In terms of hydrogen production, there is no clear set condition for maximizing it, especially regarding the initial pH (uncontrolled pH) that has been reported at values of 5.5 and 7.0. This range is way further concerning lactate (5.5 – 11.0) and caproate (5.5 – 8.5) production. In this sense, the fermentation process might be individually optimized via a careful balancing of the different operating conditions, regardless the feedstock, type of reactor and feed mode used (Table 4).

### 3.2. Co-fermentation process conditions: validation of model, significant effects, and coefficients interactions

Linear model with interaction among variables (Equation 4) was performed in order to find out the relationship between responses and process variables of co-fermentation process (Table 5). Most of the total response variation around the mean value (b0) was explained by the regression equation (Regression *p-value* significative at 5% level) and the remainder left as residual (Residual *p-value* not significance at 5% level). Furthermore, the model was found to be accurate (R^2^ of 0.72) indicating that more than 70% of the observed values could be explained by the model. The same model was used to explain the effect of variables on four alkaline pre-treatments of YMW (Ferraz-Júnior, 2020). The authors’ reported slightly higher values of R^2^ (≥0.89) than what was found in this study. This may be due to axial points not being considered in the model, indicating the variable response values were closer to the central point (i.e., greater control of casual variability) but with lower range responses.

**Table 5.** Coefficients of fitted equation (Y = b_0_ + b_1_ X_1_ + b_2_ X_2_ + b_3_ X_3_ + b_12_ X_1_ X_2_ + b_13_ X_1_ X_3_ + b_23_ X_2_ X_3_ + *e*) and its percent significance.

| Coefficients |  |
| --- | --- |
| $b_0^a$ | 0.78 |
| $b_1^a$ | 0.15 |
| $b_2$ | 0.05 |
| $b_3^a$ | 0.19 |
| $b_{12}^a$ | -0.08 |
| $b_{13}^a$ | 0.14 |
| $b_{23}$ | -0.02 |
| $e$ | $\pm 0.01$ |
| $p\text{-value}^{a, b}$ | 0.021 |
| $p\text{-value}^c$ | 0.059 |
| $R^2$ | 0.72 |

a. Significant at 5% level. b. Regression p-value. c. Residual p-value.

The factors X_1_ (YMW concentration) and X_3_ (inoculum concentration) at the levels studied are significant, indicating that they might be fixed at the lowest value when evaluated individually. Interestingly, the factor X_2_ (initial pH, *i.e*., non-controlled pH) had no influence on the co-fermentation process, suggesting that it can be fixed at any value between the two levels. Furthermore, the final pH measured from each batch reactor was between 3.4 and 5.2 regardless of its initial value, indicating that the fermentation products were able to decrease the pH even from its highest level (S. no. 15; pH of 12.7 – Table 2). This finding is also corroborated by Koyama et al., (2016) who potentially computed the use of industrial effluent in hydrogen-producing systems at its original pH (4.8). Hydrogen production under extreme conditions of pH (2.8 and 10.0) were also reported by Mota et al., (2018) and Li et al., (2020), therfore, being consistent with the current result.

Interaction between factors are important for process optimization (Ferraz Júnior et al., 2020). The individual interaction X_1_X_2_ (YMW concentration and pH) and X_1_X_3_ (concentration of YMW and inoculum) were significant. Additionally, the X_1_X_3_ interaction had the strongest effect on the process and according to this finding, both variables should be studied at their highest levels for a greater response. The best level of independent variables and their interactions on the co-fermentation process was then evaluated with a response surface plot (Figure 1).

**Figure 1.**
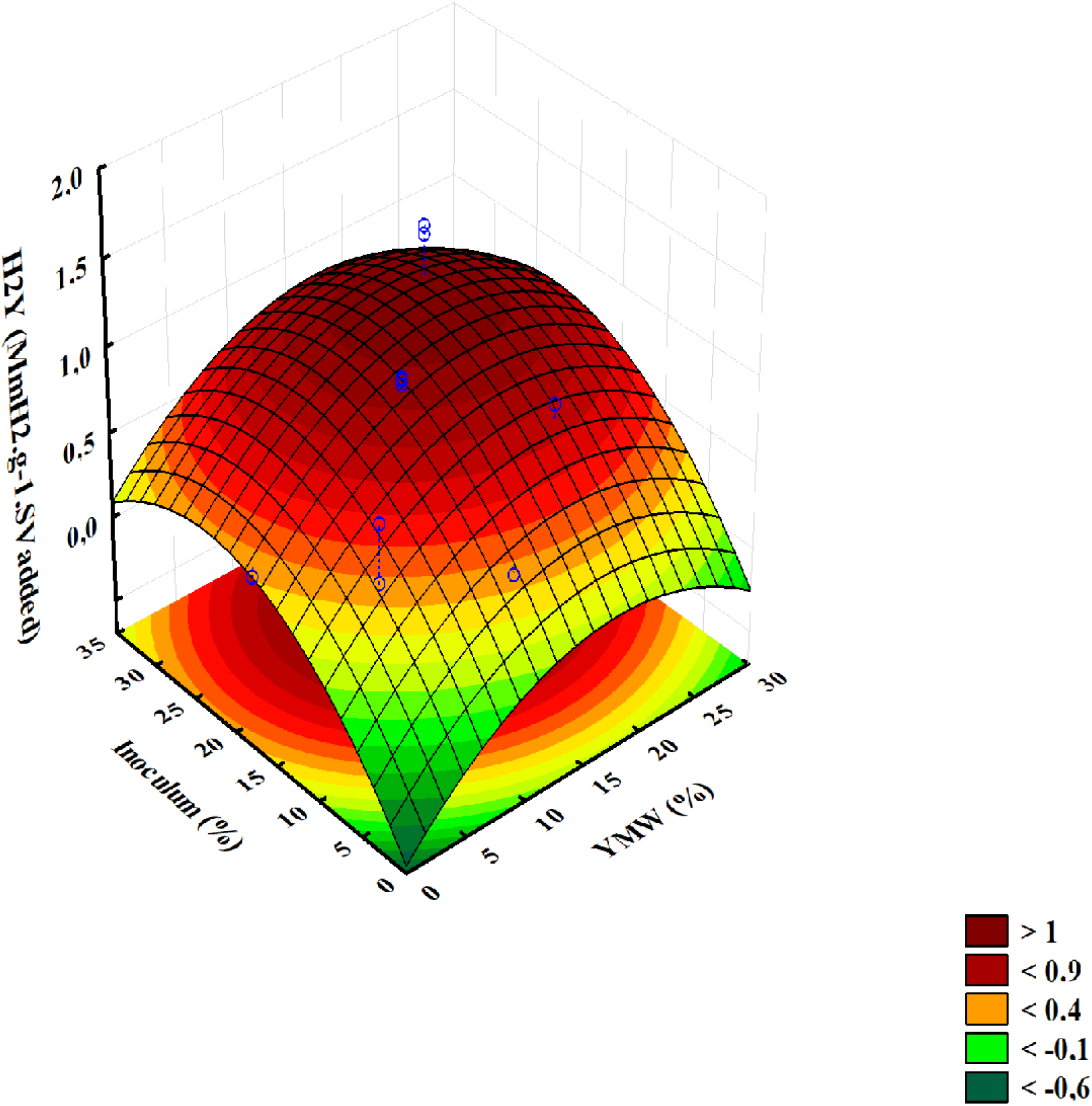
Response surface for the interactive effect on hydrogen yield (H_2_Y, MmH_2_.g^-1^ VS _added_) through co-fermentation process. Interactive effect of concentration of Yerba Mate waste and inoculum concentrations.

By applying linear regression analysis to the experimental results, Equation 11 was obtained to describe the co-fermentation process of cheese whey and Yerba Mate waste using the uncoded independent variables.

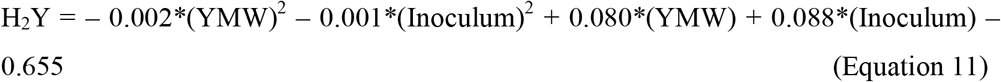

In the case, H_2_Y is the hydrogen yield in MmH_2_.g^-1^ VS _added_, YMW is the concentration of Yerba Mate in % and Inoculum is the concentration of sludge added to the reactor also in %.

### 3.4. Effects of food to microorganisms (F/M) ratios

Different ratios of F/M on hydrogen production from the co-fermentation of CW and YMW were evaluated based on the CCD experiments. The different volumes of mixtures between CW, YWM and inoculum in the batch reactors resulted in F/M ratios of 1.5, 1.8, 2.3, 2.6, 3.6, 5.6 and 9.5 g COD. g^-1^VS. The highest and lowest values of H_2_Y of co-fermentation process were archived at F/M ratio of 2.3 and 9.5 g COD. g^-1^VS, respectively, demonstrating the need for high amounts of inoculum (~ 20% w/v) able to convert complex substrates such as lignocellulosic materials in hydrogen. Similar values were observed by Nasr et al. (2011) using thin stillage as substrate. By contrast, higher ratios of F/M (10.6 – 13.3 g COD. g^-1^VS) were reported as optimum in hydrogen-producing systems (Basak et al., 2018; Ferraz Júnior et al., 2015a). The differences in the optimum F/M ratio in the literature can be attributed to the differences in the waste-type and composition as well as the anaerobic sludges.

### 3.5. Kinetic analyses of hydrogen production

Kinetics parameters can also describe the performance of processes. Modified Gompertz model was used to describe the best condition for producing hydrogen (S.no. 6; Table 2), considering the co-fermentation of CW and YMW. Concomitantly, the S.no. 12 represented the condition where the CW was used as only substrate for same purpose. To compare such behaviour between samples, it was assumed: (i) no influence of pH as previous discussed (*subhead 3.2*.) and (ii) low value for the F/M ratio (1.8 – 2.3 g COD. g^-1^VS). The Modified Gompertz model was found to describe the experimental data at an excellent level (R^2^ > 0.990) for both assays. The co-fermentation process improved the accumulated hydrogen production (AcH_2_P) and rate (R) in up to 4.5 and 7.5 folds, respectively, compared to the condition without YMW (Figure 2) (Table 6). However, the addition of YMW delayed the lag phase by 4.5 hours while, in its absence, hydrogen production occurred immediately (Table 5). It is worth mentioning that, after biogas being washed in NaOH solution (12% w/v), the hydrogen content was superior to 99% in all reactors throughout the experiment.

**Figure 2.**
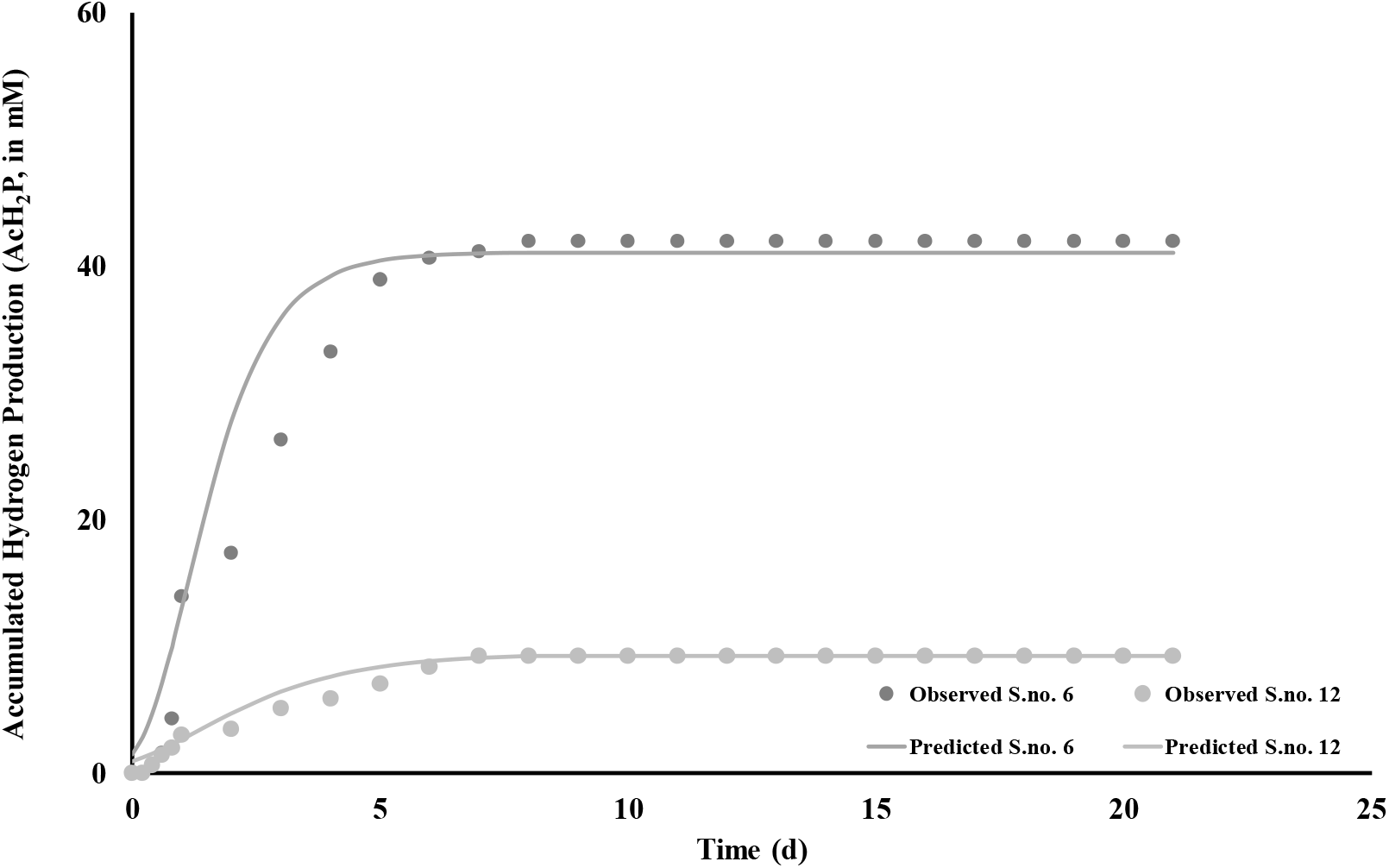
Cumulative hydrogen production (Observed) and unstructured mathematical model (Predicted) fit to the fermentative essays with (S.no. 6) and without (S.no. 12) YMW added.

**Table 6.** Kinetic parameters estimated by Modified Gompertz model from batch-reactors with and without YMW added.

| Parameters | Estimate <sup>a</sup> | Standard error <sup>a</sup> | LCL <sup>a, b</sup> | UCL <sup>a, c</sup> | R<sup>2</sup> |
| --- | --- | --- | --- | --- | --- |
| P (mM) | 41.1 / 9.4 | 1.3 / 0.5 | 38.1 / 8.4 | 44.0 / 10.5 |  |
| R (mM.d<sup>-1</sup>) | 16.2 / 2.1 | 2.2 / 0.3 | 11.3 / 1.5 | 21.1 / 2.6 | 0.992 / 0.990 |
| $\lambda$ (d) | 0.2 / -0.3 | 0.1 / 0.2 | -0.1 / -0.8 | 0.5 / 0.2 | |

a. (S.no. 6: with YMW / S.no. 12: without YMW). b. Low-confidence limit. c. Up-confidence limit.

### 3.6. Short and medium-organic acids: precursor and non-precursor of hydrogen

The total reducing sugars (TRS) in the medium source is mainly composed of glucose, lactose, xylose and arabinose, as depicted in Castelló et al., (2019) and Ferraz Júnior et al., (2020). However, the batch reactor “S. no. 12” presents only hexoses in the liquid medium, considering the absence of YM in the experiment. The TRS presented an average value of conversion of 93.4% suggesting that the organic compost provided a microbial community able to consume both pentoses and hexoses (Reactions 1-4) (Tabassum et al., 2017; Xia et al., 2015).

*Pentose conversion to acetate*

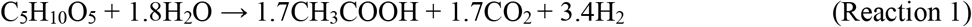

*Pentose conversion to butyrate*

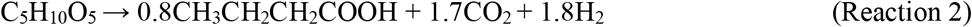

*Hexose conversion to acetate*

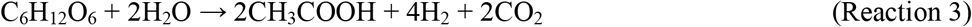

*Hexose conversion to butyrate*

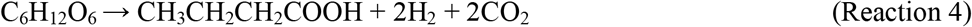

The main metabolites observed were butyrate (114-132 mM), lactate (109-140 mM) followed by acetate (16-109 mM) and ethanol (8-74 mM) (Figure 3A). As widely known, acetate-type fermentation from glucose results in 4 molH_2_.mol^-1^glucose (Reaction 5). Similarly, butyrate- and ethanol-type fermentations lead to a yield of only 2 molH_2_.mol^-1^glucose (Reaction 6-7) (Toledo-Alarcón et al., 2018).

**Figure 3.**
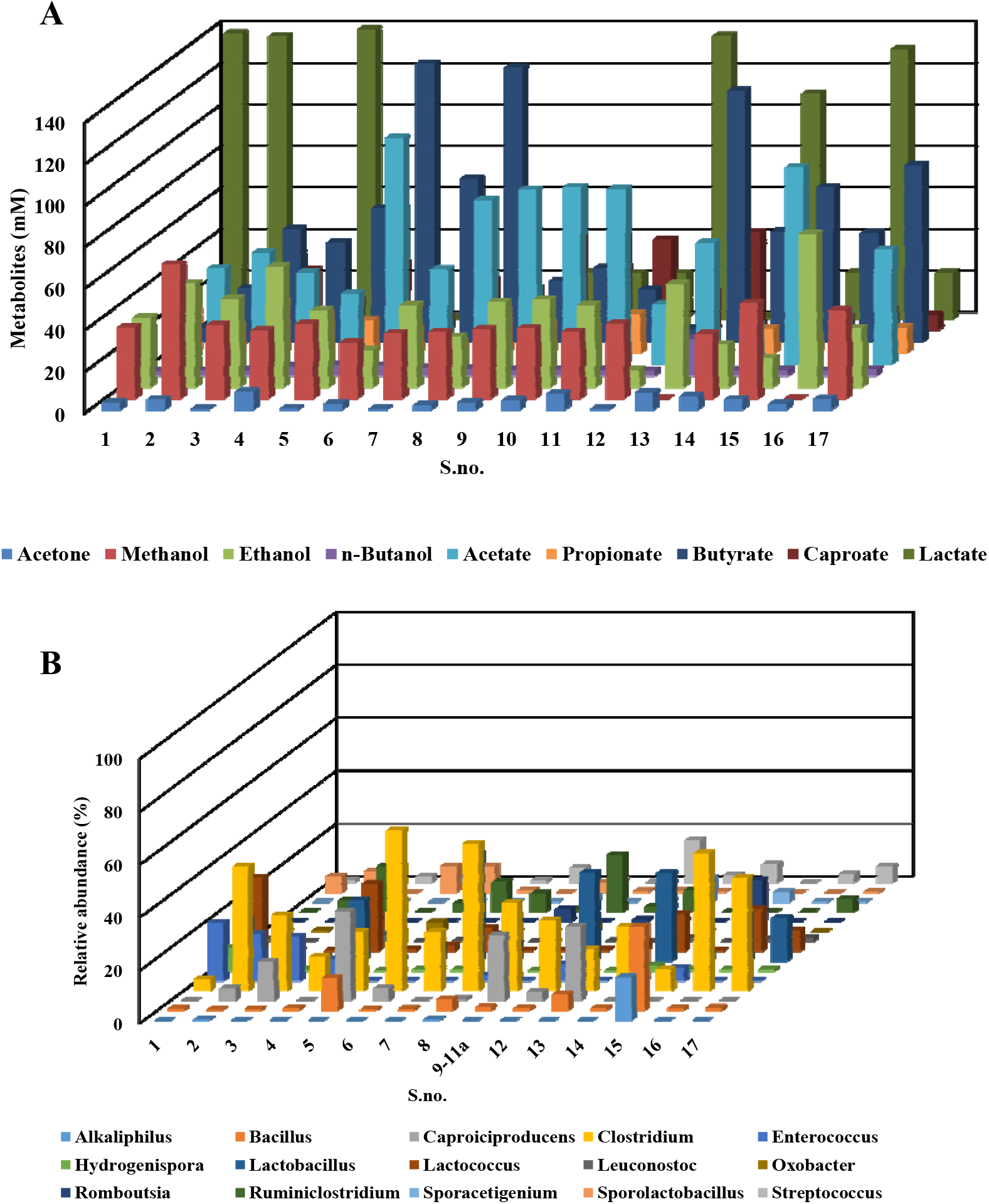
A. Metabolites (organic acids and alcohols) of CCD experiments. B. Composition of microbial community at genus level of CCD experiments. Relative abundance above 1%.

*Acetate-type fermentation*

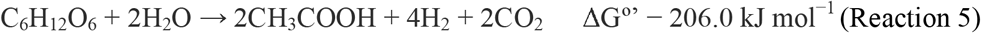

*Butyrate-type fermentation*

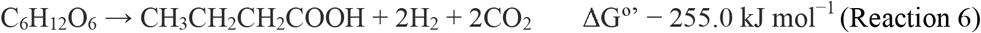

*Ethanol-type fermentation (glucose into ethanol and acetate)*

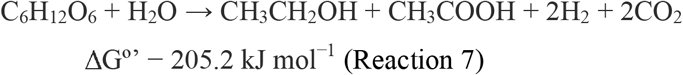

The above reactions give the impression that higher values for acetate are directly related to higher hydrogen production. However, acetate is also a product of hydrogen consumers (homoacetogens) whereas butyrate is inexorably linked to hydrogen-producing in mixed culture, and no direct hydrogen consumption pathway related to butyrate production has been reported so far (Guo et al., 2014). Furthermore, butyrate formation reaction is more energetically favorable, considering the Gibb’s free energy (Reaction 6). In turn, ethanol-type fermentation occurs in condition with high acetate concentration and low pH (lower than 4) (Mota et al., 2018).

The theoretical hydrogen production was calculated for each batch reactor. The measured hydrogen ranged between 6.3% and 41.6% of the theoretical hydrogen computed, suggesting homoacetogenesis play a key role in all batch reactors (Reaction 8). This finding is corroborated by the estimation of the measured acetate from homoacetogenesis (Equation 8). Acetate issued by such a pathway reached values up to 94.6% which explains the “low values” of hydrogen production and supports the butyrate-type fermentation as the main hydrogen-producing pathway in this study.

The increment of agitation speed might be a strategy to avoid hydrogen consumption by homoacetogens (Montiel Corona et al., 2018). These authors observed a depletion of 9% in homoacetogenesis after increasing the agitation speed. Alternatively, biogas sequestration from the headspace of a fermentative system was able to lower the availability of hydrogen in the liquid medium and, thus, minimizing homoacetogens (Ferraz Júnior et al., 2020).

*Homoacetogenesis (the Wood–Ljungdahl pathway)*

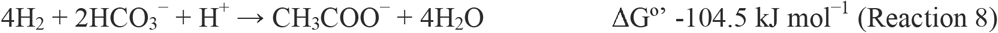

Residual sugars, hydrogen production and metabolites represented between and 70.2% and 83.8% of the COD fed to batch reactors. The organic matter conversion into biomass was not computed (Supplementary Table 1).

The experiments performed give also suitable information about the production of lactate, an added-value compound used to produce poly-lactic acid, a biodegradable plastic (Parra-Ramírez et al., 2019) and, interestingly, caproate, a medium-chain organic acid used as feed additive, plant growth promoter, etc. (Pan et al., 2020). Lactate is often reported in mesophilic hydrogen producing systems as an inhibitor of biohydrogen production process and rarely discussed as a commercial product (Reaction 9). In this study, high values of lactate (~ 140 mM) were obtained at different values of pH (5.9, 8.5 and 11.0) of co-fermentation process (Table 3A). Lactate production, separation and purification was economically viable for some of the scales evaluated at a value of 1.89 USD.kg^-1^ (Parra-Ramírez et al., 2019). Yet, lactate jointly with ethanol are reported as ideal substrates for supplying electrons during carboxylic acid chain elongation through the reverse ß-oxidation (RBO) reaction (Barker and Taha, 1942) (Reactions 10-11). In this process, the sequential formation of butyrate and, then, caproate from acetate is possible (Cavalcante et al., 2017). The maximum caproate production observed was ~ 45 mM at pH 8.5 of the CW and YMW dark fermentation (Table 3A). Its production probably occurred in two steps: (i) fermentation of organic matter and hydrogen consumption to acetate production via the *Wood–Ljungdahl* pathway followed by (ii) the RBO pathway. Furthermore, the market price of caproate is more than 10 times higher than that of ethanol (Cavalcante et al., 2017). Despite these high values of short- and medium-chain organic acids production here observed, more detailed research on this topic must be performed in order to optimize the process, considering their productivity and yield.

*Lactic-type fermentation (glucose into lactate and ethanol)*

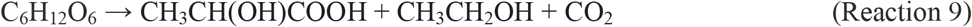

*Overall production of n-caproate from lactate*

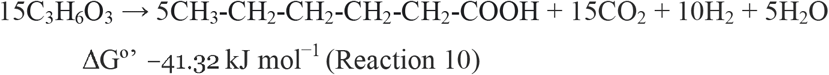

*Overall production of n-caproate from ethanol and acetate*

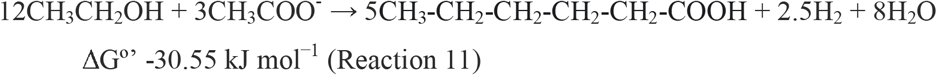

### 3.7. Taxonomic profile of the microbial community in the co-fermentation batch reactors

16S ribosomal RNA gene sequences were analyzed to characterize the microbial community structure and reveal the CW and YMW co-fermentation conditions-associated changes (Figure 3B). According to the results, the most abundant microorganisms detected in the co-fermentation process were related to the following roles: (i) hydrogen production (*Alkaliphilus, Bacillus, Clostridium, Romboutsia, Ruminiclostridium* and *Sporacetigenium*) (Ferraz Júnior et al., 2013, 2014, 2015a, 2015b, An et al., 2018, An et al., 2020, Bu et al., 2021) (ii) ethanol-hydrogen co-production (*Hydrogenispora*) (Liu et al., 2014); (iii) hydrogen consumption (*Oxobacter*) (Greening et al., 2019); (iv) lactate production (*Enterococcus, Lactobacillus, Lactococcus, Leuconostoc, Romboutsia, Sporolactobacillus* and *Streptococcus*) (Castelló et al., 2020; Ferraz Júnior et al., 2017; Fuess et al., 2018) and (v) caproate production (*Caproiciproducens*) (Kim et al., 2015). These microorganisms are consistent with fermentative systems studies and coherently related to the metabolites presented in Figure 3A and discussed in subhead 3.6.

To further understand the interaction among the indicators of batch-reactor performances, a Principal Component Analysis (PCA) was performed (Figure 4). Two principal components accounted for nearly 45% of the dataset variance. The results showed two well-defined axes or principal components (PC): PC 1 which represents the main effect of lactate and its producers’ microorganisms opposing to H_2_Y, acetate and butyrate as well as caproate and *Caproiciproducens* (PC 2). These findings reinforcing that acetate-, butyrate-, caproate- and lactate-type fermentation were the main metabolic pathways *(subhead 3.6)* observed from the CCD experiments. The results also showed a low variation of ethanol and propionate, considering the variables and their levels studied. Finally, it should be noticed that the variables YMW, pH and inoculum were computed in the PCA as supplementary elements. It means that such coordinates are predicted using only the information provided by the performed PCA on active variables/individuals.

**Figure 4.**
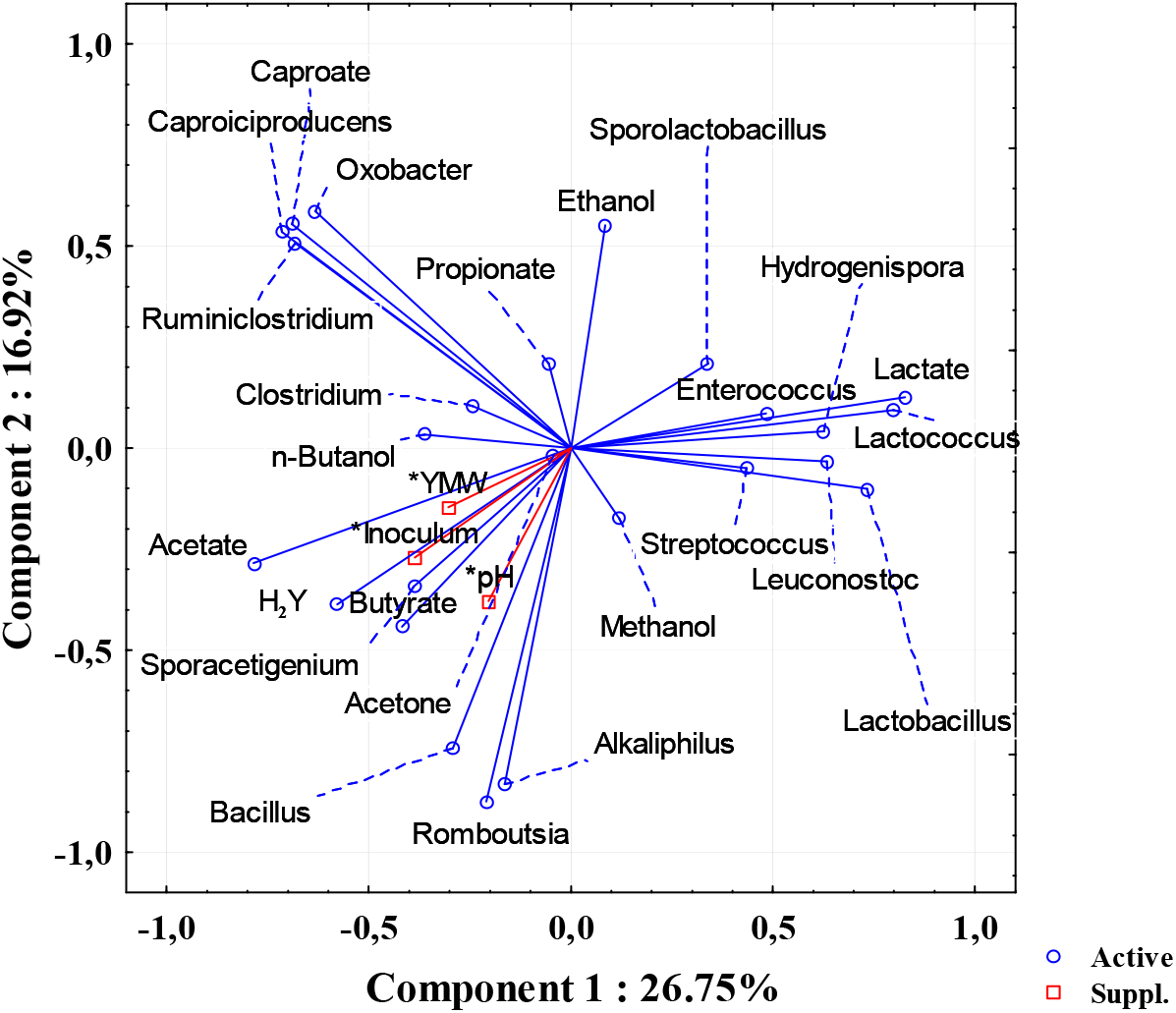
Principal components analysis (PCA) of CCD experiments.

Linked to Ferraz Júnior et al. (2014b), Ferraz Júnior et al. (2014b) and Luo et al. (Luo et al., 2011).

## 4. Conclusions

Co-fermentation of cheese whey and alkaline-pretreated Yerba Mate waste can be potentially used to produce hydrogen and short- and medium-chain organic acids. In terms of hydrogen production, the increase of inoculum and YMW concentrations had positive effects in the process while the initial pH had no significant effect on it, considering the conditions evaluated. Butyrate-type fermentation was the main hydrogen-producing pathway. Acetate from homoacetogenesis was accounted for all conditions evaluated. The Central Composite Design also indicated operating conditions to produce moderate-to-high concentrations of added value compounds, for instance, butyrate, lactate and caproate. 16S ribosomal DNA gene sequences analysis revealed five groups of microorganisms related to hydrogen, lactate and caproate production, ethanol-hydrogen co-production and hydrogen consumption. Principal Component Analysis computed three well-defined groups related to the hydrogen, lactate and caproate production.

## List of abbreviations

b_1_: linear coefficients for the YMW concentration
b_2_: linear coefficients for pH
b_3_: linear coefficients for the inoculum concentration
CCD: Central Composite Designs
CW: Cheese Whey
DNS: 3,5-Dinitrosalicylic acid method
DoE: Design of Experiments
F/M: Food-to-Microorganisms ratio
H_2_Y: Hydrogen Yield
HTS: High-Throughput Sequencing
PCA: Principal Component Analysis
RBO: Reverse β-Oxidation
RSM: Response Surface Methodology
SCOD: Soluble Chemical Demand of Oxygen
TCOD: Total Chemical Demand of Oxygen
TRS: Total Reducing Sugars
TS: Total solids
TVS: Total Volatile Solids
X_1_: YMW concentration
X_12_: coefficients for the interaction between YMW concentration and pH
X_13_: coefficients for the interaction between YMW concentration and inoculum concentration
X_2_: pH
X_23_: coefficients for the interaction between pH and inoculum concentration
X_3_: inoculum concentration
Y: Experimental Response
Ŷ: Predicted Response
YMW: Yerba Mate Waste

## Acknowledgements

The authors gratefully acknowledge support of the Clemente Estable Institute of Biological Research (IIBCE), the Program of Basic Science Development of Uruguay (PEDECIBA-Chemistry), the Uruguayan National System of Researchers and National Agency for Research and Innovation (SNI-ANII).

---

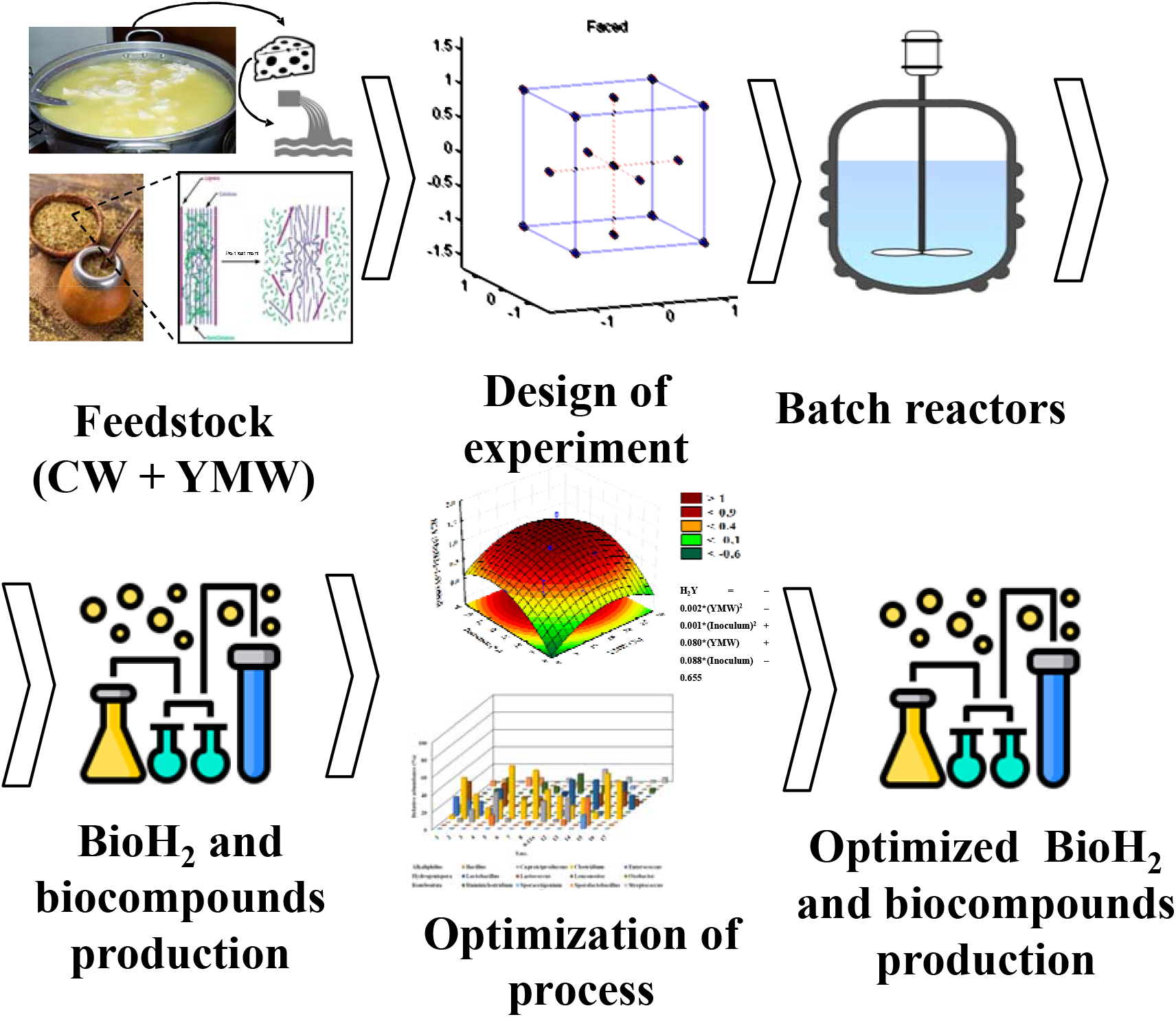

## References

Adorno, M.A.T., Hirasawa, J.S., Varesche, M.B.A., 2014. Development and Validation of Two Methods to Quantify Volatile Acids (C2-C6) by GC/FID: Headspace (Automatic and Manual) and Liquid-Liquid Extraction (LLE). Am. J. Anal. Chem. 05, 406–414. https://doi.org/10.4236/ajac.2014.57049

Akhlaghi, M., Boni, M.R., De Gioannis, G., Muntoni, A., Polettini, A., Pomi, R., Rossi, A., Spiga, D., 2017. A parametric response surface study of fermentative hydrogen production from cheese whey. Bioresour. Technol. 244, 473–483. https://doi.org/10.1016/j.biortech.2017.07.158

An, Q., Cheng, J.-R., Wang, Y.-T., Zhu, M.-J., 2020. Performance and energy recovery of single and two stage biogas production from paper sludge: Clostridium thermocellum augmentation and microbial community analysis. Renew. Energy 148, 214–222. https://doi.org/10.1016/j.renene.2019.11.142

An, Q., Wang, J.-L., Wang, Y.-T., Lin, Z.-L., Zhu, M.-J., 2018. Investigation on hydrogen production from paper sludge without inoculation and its enhancement by Clostridium thermocellum. Bioresour. Technol. 263, 120–127. https://doi.org/10.1016/j.biortech.2018.04.105

APHA, 2005. Standard methods for the examination of water and wastewater, 21ts ed. American Public Health Association, Washington, DC, New York: American Public Health Association.

Barker, H.A., Taha, S.M., 1942. Clostridium kluyverii, an Organism Concerned in the Formation of Caproic Acid from Ethyl Alcohol. J. Bacteriol. 43, 347–63. https://doi.org/10.1128/JB.43.3.347-363.1942

Basak, B., Fatima, A., Jeon, B.-H., Ganguly, A., Chatterjee, P.K., Dey, A., 2018. Process kinetic studies of biohydrogen production by co-fermentation of fruit-vegetable wastes and cottage cheese whey. Energy Sustain. Dev. 47, 39–52. https://doi.org/10.1016/j.esd.2018.08.004

Bina, B., Amin, M.M., Pourzamani, H., Fatehizadeh, A., Ghasemian, M., Mahdavi, M., Taheri, E., 2019. Biohydrogen production from alkaline wastewater: The stoichiometric reactions, modeling, and electron equivalent. MethodsX 6, 1496–1505. https://doi.org/10.1016/j.mex.2019.06.013

Borin, G.P., Alves, R.F., Ferraz Júnior, A.D.N., 2019. Current Status of Biotechnological Processes in the Biofuel Industries, in: Bioprocessing for Biomolecules Production. John Wiley & Sons, Ltd, Chichester, UK, pp. 47–69. https://doi.org/10.1002/9781119434436.ch3

Borshchevskaya, L.N., Gordeeva, T.L., Kalinina, A.N., Sineokii, S.P., 2016. Spectrophotometric determination of lactic acid. J. Anal. Chem. 71, 755–758. https://doi.org/10.1134/S1061934816080037

Box G.E, Wilson, K.., 1951. On the Experimental Attainment of Optimum Conditions. J. R. Stat. Soc. 13, 1–45.

Bu, J., Wang, Y.-T., Deng, M.-C., Zhu, M.-J., 2021. Enhanced enzymatic hydrolysis and hydrogen production of sugarcane bagasse pretreated by peroxyformic acid. Bioresour. Technol. 326, 124751. https://doi.org/10.1016/j.biortech.2021.124751

Caporaso, J.G., Lauber, C.L., Walters, W. a, Berg-Lyons, D., Huntley, J., Fierer, N., Owens, S.M., Betley, J., Fraser, L., Bauer, M., Gormley, N., Gilbert, J. a, Smith, G., Knight, R., 2012. Ultra-high-throughput microbial community analysis on the Illumina HiSeq and MiSeq platforms. ISME J. 6, 1621–1624. https://doi.org/10.1038/ismej.2012.8

Caporaso, J.G., Lauber, C.L., Walters, W.A., Berg-Lyons, D., Lozupone, C.A., Turnbaugh, P.J., Fierer, N., Knight, R., 2011. Global patterns of 16S rRNA diversity at a depth of millions of sequences per sample. Proc. Natl. Acad. Sci. U. S. A. 108 Suppl, 4516–22. https://doi.org/10.1073/pnas.1000080107

Castelló, E., Nunes Ferraz-Júnior, A.D., Andreani, C., Anzola-Rojas, M. del P., Borzacconi, L., Buitrón, G., Carrillo-Reyes, J., Gomes, S.D., Maintinguer, S.I., Moreno-Andrade, I., Palomo-Briones, R., Razo-Flores, E., Schiappacasse-Dasati, M., Tapia-Venegas, E., Valdez-Vázquez, I., Vesga-Baron, A., Zaiat, M., Etchebehere, C., 2020. Stability problems in the hydrogen production by dark fermentation: Possible causes and solutions. Renew. Sustain. Energy Rev. 119, 109602. https://doi.org/10.1016/j.rser.2019.109602

Cavalcante, W. de A., Leitão, R.C., Gehring, T.A., Angenent, L.T., Santaella, S.T., 2017. Anaerobic fermentation for n-caproic acid production: A review. Process Biochem. 54, 106–119. https://doi.org/10.1016/j.procbio.2016.12.024

Chen, S., Song, L., Dong, X., 2006. Sporacetigenium mesophilum gen. nov., sp. nov., isolated from an anaerobic digester treating municipal solid waste and sewage. Int. J. Syst. Evol. Microbiol. 56, 721–725. https://doi.org/10.1099/ijs.0.63686-0

Claesson, M.J., O’Sullivan, O., Wang, Q., Nikkilä, J., Marchesi, J.R., Smidt, H., de Vos, W.M., Ross, R.P., O’Toole, P.W., 2009. Comparative Analysis of Pyrosequencing and a Phylogenetic Microarray for Exploring Microbial Community Structures in the Human Distal Intestine. PLoS One 4, e6669. https://doi.org/10.1371/journal.pone.0006669

Dareioti, M.A., Vavouraki, A.I., Kornaros, M., 2014. Effect of pH on the anaerobic acidogenesis of agroindustrial wastewaters for maximization of bio-hydrogen production: A lab-scale evaluation using batch tests. Bioresour. Technol. 162, 218–227. https://doi.org/10.1016/j.biortech.2014.03.149

Edgar, R.C., Haas, B.J., Clemente, J.C., Quince, C., Knight, R., 2011. UCHIME improves sensitivity and speed of chimera detection. Bioinformatics 27, 2194–2200. https://doi.org/10.1093/bioinformatics/btr381

Fernández, C., Carracedo, B., Martínez, E.J., Gómez, X., Morán, A., 2014. Application of a packed bed reactor for the production of hydrogen from cheese whey permeate: Effect of organic loading rate. J. Environ. Sci. Heal. Part A 49, 210–217. https://doi.org/10.1080/10934529.2013.838885

Ferraz-Júnior; Antônio Djalma Nunes, Etchelet, M.I., Braga, A.F.M., Clavijo, L., Loaces, I., Noya, F., Etchebehere, C., 2020. Alkaline pretreatment of yerba mate (Ilex paraguariensis) waste for unlocking low-cost cellulosic biofuel. Fuel 266, 117068. https://doi.org/10.1016/j.fuel.2020.117068

Ferraz Júnior, A.D.N., 2013. Digestão anaeróbia da vinhaça da cana de açúcar em reator acidogênico de leito fixo seguido de reator metanogênico de manta de lodo. Universidade de São Paulo, São Carlos. https://doi.org/10.11606/T.18.2013.tde-27082014-092345

Ferraz Júnior, A.D.N., Damásio, A.R.L., Paixão, D.A.A., Alvarez, T.M., Squina, F.M., 2017. Applied Metagenomics for Biofuel Development and Environmental Sustainability, in: Advances of Basic Science for Second Generation Bioethanol from Sugarcane. Springer International Publishing, Cham, pp. 107–129. https://doi.org/10.1007/978-3-319-49826-3_7

Ferraz Júnior, A.D.N., Etchebehere, C., Zaiat, M., 2015a. High organic loading rate on thermophilic hydrogen production and metagenomic study at an anaerobic packed-bed reactor treating a residual liquid stream of a Brazilian biorefinery. Bioresour. Technol. 186, 81–88. https://doi.org/10.1016/j.biortech.2015.03.035

Ferraz Júnior, A.D.N., Etchebehere, C., Zaiat, M., 2015b. Mesophilic hydrogen production in acidogenic packed-bed reactors (apbr) using raw sugarcane vinasse as substrate: Influence of support materials. Anaerobe. https://doi.org/10.1016/j.anaerobe.2015.04.008

Ferraz Júnior, A.D.N., Pages, C., Latrille, E., Bernet, N., Zaiat, M., Trably, E., 2020. Biogas sequestration from the headspace of a fermentative system enhances hydrogen production rate and yield. Int. J. Hydrogen Energy. https://doi.org/10.1016/j.ijhydene.2020.02.064

Ferraz Júnior, A.D.N., Wenzel, J., Etchebehere, C., Zaiat, M., 2014a. Effect of organic loading rate on hydrogen production from sugarcane vinasse in thermophilic acidogenic packed bed reactors. Int. J. Hydrogen Energy 39, 16852–16862. https://doi.org/10.1016/j.ijhydene.2014.08.017

Ferraz Júnior, A.D.N., Zaiat, M., Gupta, M., Elbeshbishy, E., Hafez, H., Nakhla, G., 2014b. Impact of organic loading rate on biohydrogen production in an up-flow anaerobic packed bed reactor (UAnPBR). Bioresour. Technol. 164, 371–9. https://doi.org/10.1016/j.biortech.2014.05.011

Fuess, L.T., Ferraz, A.D.N., Machado, C.B., Zaiat, M., 2018. Temporal dynamics and metabolic correlation between lactate-producing and hydrogen-producing bacteria in sugarcane vinasse dark fermentation: The key role of lactate. Bioresour. Technol. 247, 426–433. https://doi.org/10.1016/j.biortech.2017.09.121

Greening, C., Geier, R., Wang, C., Woods, L.C., Morales, S.E., McDonald, M.J., Rushton-Green, R., Morgan, X.C., Koike, S., Leahy, S.C., Kelly, W.J., Cann, I., Attwood, G.T., Cook, G.M., Mackie, R.I., 2019. Diverse hydrogen production and consumption pathways influence methane production in ruminants. ISME J. 13, 2617–2632. https://doi.org/10.1038/s41396-019-0464-2

Grosser, A., Neczaj, E., 2018. Sewage sludge and fat rich materials co-digestion - Performance and energy potential. J. Clean. Prod. 198, 1076–1089. https://doi.org/10.1016/j.jclepro.2018.07.124

Guo, X.M., Trably, E., Latrille, E., Carrere, H., Steyer, J.-P., 2014. Predictive and explicative models of fermentative hydrogen production from solid organic waste: Role of butyrate and lactate pathways. Int. J. Hydrogen Energy 39, 7476–7485. https://doi.org/10.1016/j.ijhydene.2013.08.079

Kim, B.-C., Seung Jeon, B., Kim, S., Kim, H., Um, Y., Sang, B.-I., 2015. Caproiciproducens galactitolivorans gen. nov., sp. nov., a bacterium capable of producing caproic acid from galactitol, isolated from a wastewater treatment plant. Int. J. Syst. Evol. Microbiol. 65, 4902–4908. https://doi.org/10.1099/ijsem.0.000665

Kim, D.-H., Kim, S.-H., Jung, K.-W., Kim, M.-S., Shin, H.-S., 2011. Effect of initial pH independent of operational pH on hydrogen fermentation of food waste. Bioresour. Technol. 102, 8646–8652. https://doi.org/10.1016/j.biortech.2011.03.030

Koyama, M.H., Messias, M., Júnior, A., Zaiat, M., 2016. Kinetics of thermophilic acidogenesis of typical Brazilian sugarcane vinasse. Energy 116, 1097–1103. https://doi.org/10.1016/j.energy.2016.10.043

Lee, Z.-K., Li, S.-L., Lin, J.-S., Wang, Y.-H., Kuo, P.-C., Cheng, S.-S., 2008. Effect of pH in fermentation of vegetable kitchen wastes on hydrogen production under a thermophilic condition. Int. J. Hydrogen Energy 33, 5234–5241. https://doi.org/10.1016/j.ijhydene.2008.05.006

Levin, D.B., Pitt, L., Love, M., 2004. Biohydrogen production: prospects and limitations to practical application 29, 173–185. https://doi.org/10.1016/S0360-3199(03)00094-6

Li, X., Guo, L., Liu, Y., Wang, Y., She, Z., Gao, M., Zhao, Y., 2020. Effect of salinity and pH on dark fermentation with thermophilic bacteria pretreated swine wastewater. J. Environ. Manage. 271, 111023. https://doi.org/10.1016/j.jenvman.2020.111023

Liu, Y., 1996. Bioenergetic interpretation on the S0X0 ratio in substrate-sufficient batch culture. Water Res. 30, 2766–2770. https://doi.org/10.1016/S0043-1354(96)00157-1

Liu, Y., Qiao, J.-T., Yuan, X.-Z., Guo, R.-B., Qiu, Y.-L., 2014. Hydrogenispora ethanolica gen. nov., sp. nov., an anaerobic carbohydrate-fermenting bacterium from anaerobic sludge. Int. J. Syst. Evol. Microbiol. 64, 1756–1762. https://doi.org/10.1099/ijs.0.060186-0

Lovato, G., Albanez, R., Stracieri, L., Ruggero, L.S., Ratusznei, S.M., Rodrigues, J.A.D., 2018. Hydrogen production by co-digesting cheese whey and glycerin in an AnSBBR: Temperature effect. Biochem. Eng. J. 138, 81–90. https://doi.org/10.1016/j.bej.2018.07.007

Lovato, G., Augusto, I.M.G., Ferraz Júnior, A.D.N., Albanez, R., Ratusznei, S.M., Etchebehere, C., Zaiat, M., Rodrigues, J.A.D., 2021. Reactor start-up strategy as key for high and stable hydrogen production from cheese whey thermophilic dark fermentation. Int. J. Hydrogen Energy. https://doi.org/10.1016/j.ijhydene.2021.06.010

Lucas, S.D.M., Peixoto, G., Mockaitis, G., Zaiat, M., Gomes, S.D., 2015. Energy recovery from agro-industrial wastewaters through biohydrogen production: Kinetic evaluation and technological feasibility. Renew. Energy 75, 496–504. https://doi.org/10.1016/j.renene.2014.10.025

Luo, G., Karakashev, D., Xie, L., Zhou, Q., Angelidaki, I., 2011. Long-term effect of inoculum pretreatment on fermentative hydrogen production by repeated batch cultivations: Homoacetogenesis and methanogenesis as competitors to hydrogen production. Biotechnol. Bioeng. 108, 1816–1827. https://doi.org/10.1002/bit.23122

Luongo, V., Policastro, G., Ghimire, A., Pirozzi, F., Fabbricino, M., 2019. Repeated-Batch Fermentation of Cheese Whey for Semi-Continuous Lactic Acid Production Using Mixed Cultures at Uncontrolled pH. Sustainability 11, 3330. https://doi.org/10.3390/su11123330

Marone, A., Varrone, C., Fiocchetti, F., Giussani, B., Izzo, G., Mentuccia, L., Rosa, S., Signorini, A., 2015. Optimization of substrate composition for biohydrogen production from buffalo slurry co-fermented with cheese whey and crude glycerol, using microbial mixed culture. Int. J. Hydrogen Energy 40, 209–218. https://doi.org/10.1016/j.ijhydene.2014.11.008

Miller, G.L., 1959. Use of Dinitrosalicylic Acid Reagent for Determination of Reducing Sugar. Anal. Chem. 31, 426–428. https://doi.org/10.1021/ac60147a030

Mohd Yasin, N.H., Rahman, N.A., Man, H.C., Mohd Yusoff, M.Z., Hassan, M.A., 2011. Microbial characterization of hydrogen-producing bacteria in fermented food waste at different pH values. Int. J. Hydrogen Energy 36, 9571–9580. https://doi.org/10.1016/j.ijhydene.2011.05.048

Montiel Corona, V., Razo-Flores, E., Corona, Elías; Razo-Flores, V., 2018. Continuous hydrogen and methane production from Agave tequilana bagasse hydrolysate by sequential proce ss to maximize energy recovery efficiency. Bioresour. Technol. 249, 334–341. https://doi.org/10.1016/j.biortech.2017.10.032

Mota, V.T., Ferraz Júnior, A.D.N., Trably, E., Zaiat, M., 2018. Biohydrogen production at pH below 3.0: Is it possible? Water Res. 128. https://doi.org/10.1016/j.watres.2017.10.060

Pan, X.-R., Huang, L., Fu, X.-Z., Yuan, Y.-R., Liu, H.-Q., Li, W.-W., Yu, L., Zhao, Q.-B., Zuo, J., Chen, L., Lam, P.K.-S., 2020. Long-term, selective production of caproate in an anaerobic membrane bioreactor. Bioresour. Technol. 302, 122865. https://doi.org/10.1016/j.biortech.2020.122865

Parra-Ramírez, D., Martinez, A., Cardona, C.A., 2019. Lactic acid production from glucose and xylose using the lactogenic Escherichia coli strain JU15: Experiments and techno-economic results. Bioresour. Technol. 273, 86–92. https://doi.org/10.1016/j.biortech.2018.10.061

Rao, R., Basak, N., 2021. Optimization and modelling of dark fermentative hydrogen production from cheese whey by Enterobacter aerogenes 2822. Int. J. Hydrogen Energy 46, 1777–1800. https://doi.org/10.1016/j.ijhydene.2020.10.142

Sun, C., Xia, A., Liao, Q., Fu, Q., Huang, Y., Zhu, X., Wei, P., Lin, R., Murphy, J.D., 2018. Improving production of volatile fatty acids and hydrogen from microalgae and rice residue: Effects of physicochemical characteristics and mix ratios. Appl. Energy 230, 1082–1092. https://doi.org/10.1016/j.apenergy.2018.09.066

Tabassum, M.R., Xia, A., Murphy, J.D., 2017. Potential of seaweed as a feedstock for renewable gaseous fuel production in Ireland. Renew. Sustain. Energy Rev. 68, 136–146. https://doi.org/10.1016/j.rser.2016.09.111

Toledo-Alarcón, J., Capson-Tojo, G., Marone, A., Paillet, F., Ferraz Júnior, A.D.N., Chatellard, L., Bernet, N., Trably, E., 2018. Basics of bio-hydrogen production by dark fermentation, Green Energy and Technology. https://doi.org/10.1007/978-981-10-7677-0_6

Xia, A., Cheng, J., Ding, L., Lin, R., Song, W., Su, H., Zhou, J., Cen, K., 2015. Substrate consumption and hydrogen production via co-fermentation of monomers derived from carbohydrates and proteins in biomass wastes. Appl. Energy 139, 9–16. https://doi.org/10.1016/j.apenergy.2014.11.016

Xia, A., Jacob, A., Tabassum, M.R., Herrmann, C., Murphy, J.D., 2016. Production of hydrogen, ethanol and volatile fatty acids through co-fermentation of macro- and micro-algae. Bioresour. Technol. 205, 118–125. https://doi.org/10.1016/j.biortech.2016.01.025

Yang, G., Hu, Y., Wang, J., 2019. Biohydrogen production from co-fermentation of fallen leaves and sewage sludge. Bioresour. Technol. 285, 121342. https://doi.org/10.1016/j.biortech.2019.121342

Yin, Y., Chen, Y., Wang, J., 2021. Co-fermentation of sewage sludge and algae and Fe2+ addition for enhancing hydrogen production. Int. J. Hydrogen Energy 46, 8950–8960. https://doi.org/10.1016/j.ijhydene.2021.01.009

