## Supplementary material - Table 1 for "Potentialities of biotechnological recovery of hydrogen and short- and medium-chain organic acids from the co-fermentation of cheese whey and Yerba Mate (*Ilex paraguariensis*) waste"

Table S1. Estimation of theoretical hydrogen production, recuperation of COD fed and acetate from homoacetogenesis at the end of CCD experiments.

| S.no. | Experimental H_2_ (mM) | Theoretical H_2_ (mM)^a^ | Exp. H_2_/Theoretical H_2_ (%) | COD rec. (%)^b^ | Acetate from homoacetogenesis (mM)^c^ | Acetate from homoacetogenesis/Total acetatec (%) |
| --- | --- | --- | --- | --- | --- | --- |
| 1 | 11.2 | 26.8 | 41.6 | 79.9 | 2.6 | 16.1 |
| 2 | 18.0 | 131.7 | 13.7 | 70.5 | 18.9 | 40.9 |
| 3 | 22.3 | 192.0 | 11.6 | 83.6 | 28.3 | 52.5 |
| 4 | 19.5 | 165.0 | 11.8 | 76.6 | 24.3 | 54.8 |
| 5 | 18.0 | 181.2 | 9.9 | 83.3 | 27.2 | 79.5 |
| 6 | 41.9 | 475.0 | 8.8 | 72.5 | 72.2 | 66.1 |
| 7 | 26.0 | 243.4 | 10.7 | 83.6 | 36.2 | 79.1 |
| 8 | 40.3 | 402.5 | 10.0 | 75.2 | 60.4 | 76.2 |
| 9 | 29.5 | 213.2 | 13.8 | 70.5 | 30.6 | 36.3 |
| 10 | 27.0 | 229.8 | 11.7 | 73.1 | 33.8 | 39.6 |
| 11 | 27.3 | 200.7 | 13.6 | 70.5 | 28.9 | 34.1 |
| 12 | 9.6 | 70.6 | 13.6 | 83.9 | 10.2 | 35.2 |
| 13 | 22.6 | 358.9 | 6.3 | 70.5 | 56.0 | 95.6 |
| 14 | 30.4 | 132.6 | 22.9 | 70.2 | 17.0 | 89.7 |
| 15 | 30.7 | 330.4 | 9.3 | 70.3 | 50.0 | 52.6 |
| 16 | 10.2 | 156.2 | 6.5 | 76.5 | 24.3 | 94.6 |
| 17 | 25.1 | 268.4 | 9.4 | 72.1 | 40.5 | 73.3 |

a. Theoretical hydrogen production and yield are based on the acetate, butyrate and propionate produced according to Ferraz Júnior et al. (2014b). b. Calculated according to Ferraz Júnior et al. (2014b). c. Acetate from homoacetogenesis was calculated according to Luo et al. (Luo et al., 2011).
